# Long noncoding RNA Hottip maintained skeletal homeostasis *via* suppressing the enhancer of zeste homolog 2 (Ezh2)/histone methylation regulatory axis

**DOI:** 10.1101/2023.11.13.566872

**Authors:** Zhi-Peng Li, Yong-Xin Mai, Shu-Ting Zhou, Chuan-jian Shi, Jiang Shao, Puping Liang, Wei-cheng Liang, Jin-fang Zhang

## Abstract

Recent evidence suggests that long noncoding RNAs (lncRNAs) play a crucial role in regulating bone remodeling and skeletal homeostasis by coordinating the development of osteoblasts and osteoclasts. Several lncRNAs have been identified to participate in bone formation and resorption processes. Among them, the well-known oncogenic lncRNA, Hottip, has been reported to be involved in osteogenesis regulation. However, the specific function and underlying mechanisms remain poorly understood.

In this study, we investigated the role of lncRNA Hottip in bone remodeling and skeletal homeostasis. Hottip knockout mice exhibited disrupted bone metabolism, abnormal bone tissue, and compromised bone quality, resulting in delayed fracture healing. In vitro experiments demonstrated that Hottip knockdown inhibited osteoblast differentiation while promoting osteoclast differentiation, with the opposite effect observed upon Hottip overexpression. Mechanistically, Hottip physically interacted with EZH2, leading to its degradation and facilitating the transcription of osteogenic genes by suppressing H3K9me3 and H3K27me3. In vivo experiments further validated the potential of Hottip overexpression to promote bone regeneration and accelerate fracture healing.

In conclusion, our study reveals Hottip as a critical regulator in the differentiation of osteoblasts and osteoclasts, crucial for maintaining skeletal homeostasis. This lncRNA shows potential as a promising therapeutic target for bone regeneration.

## Introduction

Bone continuously remodels and regenerates throughout life^1^. The equilibrium between bone formation and bone resorption is the key to maintain bone homeostasis^2^. Osteoblasts, derived from mesenchymal stem cells, were responsible for bone formation including the production of bone matrix and mineralization^3^. On the contrary, osteoclasts are the primary bone resorptive cells and originate from precursors of the hematopoietic lineage^4^. Pre-osteoclasts differentiate into multinucleated giant cells that degrade bone tissue. Maintaining a balanced interaction between osteoblasts and osteoclasts is crucial for adjusting bone resorption and formation^5^. Once it is disrupted, the bone remodeling process is disturbed and regeneration is damaged, which leads to bone-related diseases such as osteoporosis, fracture nonunion, etc.

Long noncoding RNAs (lncRNAs) have emerged as crucial regulators in diverse biological processes and disease pathogenesis. Several lncRNAs, including H19, Malat1, Tug1, and Hotair, play significant roles in bone formation.^6–9^. On the other hand, several other lncRNAs such as AK077216, MIRG and LINC00311 were demonstrated to participate in osteoclastogenesis and bone resorption^10–12^. LncRNAs may regulate the balance between osteoblast and osteoclast differentiation, impacting bone remodeling and regeneration. Hottip, a 3.7k nucleotide long non-coding RNA (lncRNA), is linked to malignancies and affects cancer patient outcomes. Its role in other diseases is not well-known. Hottip knockout (KO) mice showed muscle defects, hindlimb skeletal malformations, and shorter calcaneum bones, suggesting its potential role in musculoskeletal development^13^. Further investigation also demonstrated that Hottip promoted osteogenesis of mesenchymal stem cells (MSCs)^14^. We thereby speculate that Hottip may participate in skeletal development through mediating bone remodeling.

In this study, Hottip KO mice were generated using the CRISPR/Cas9 system, revealing imbalanced bone homeostasis and impaired bone quality. Delayed bone fracture healing was also observed in the KO mice. Hottip was found to promote osteogenic differentiation of MSCs while suppressing osteoclast differentiation of MNCs, suggesting its role as a regulator in maintaining bone formation and resorption balance. Mechanistically, Hottip recruited and potentiated EZH2-CDK1 interaction, leading to the disruption of EZH2-catalyzed histone H3K9 and K27 trimethylation. This novel epigenetic mechanism demonstrates Hottip’s role in bone regeneration, presenting potential clinical applications.

## Materials and methods

A comprehensive elucidation of the experimental methodologies employed is detailed within the supplementary materials and methods section.

### Knockout mice

Hottip KO mice were generated using the CRISPR/Cas9 system from Cyagen Biosciences (Guangzhou, China). The targeting vector containing gRNA sequences (gRNA1:CTGGATTGGACCAGGTACAACGG,gRNA2:ATTTCTATGAGATGTCTA TAGGG) was constructed. Positive clones were microinjected into C57BL/6J mouse blastocysts, yielding three positive F0 mice identified through PCR examination. F1 mice were obtained by backcrossing with WT mice and identified using PCR examination. All animals were housed in a specific pathogen-free environment at a constant temperature of 20-24°C with free access to food and water. Ethical guidelines of Guangzhou University of Chinese Medicine were followed for animal care and use.

### Metabonomics and transcriptomics

The metabolomics and transcriptomics were performed by Novogene Co. Ltd. and Shanghai luming biological technology Co. Ltd, respectively.The differently expressed metabolites and genes were analyzed and then heat map and volcano plots were performed using R3.6.3. Gene set enrichment analysis (GSEA) was conducted to identify the potential candidates and signaling.

### MSCs isolation and osteogenic differentiation

Mouse MSCs were isolated from the tibia and femoral marrow compartments of 12-week-old KO and WT mice by flushing the bone marrow into a 10-cm cell culture dish. The non-adherent cells were removed by changing the medium, and MSCs were characterized by flow cytometry after 72 hours ^15^. For osteogenic differentiation, MSCs were incubated with the classical cocktails containing dexamethasone, ascorbic acid, and glycerol 2-phosphate^16^.The differentiation medium was replaced every three days. MSCs were stocked for future usage.

### Mononuclear cells (MNCs) and osteoclast differentiation

Primary MNCs were isolated from femoral marrow compartments with α-minimal essential medium (α-MEM)^17^. The bone marrow cells were incubated in α-MEM medium supplemented with 10% FBS. The unattached cells were collected and plated in 96-well plates with MCSF (50ng/ml, PeproTech, Rocky hill, USA)^18^added. Osteoclast differentiation was induced by adding 25ng/ml MCSF and 50ng/ml RANKL (R&D Systems, Inc., Minneapolis, MN, USA).

### Plasmids construction and establishment of stable cells

Hottip knockdown (shHottip) and Hottip overexpression (pHottip) lentivirus plasmids were purchased from VectorBuilder (Yunzhou Biotechnology, Guangzhou, China). These plasmids were identified by DNA sequencing. The production and purification of the lentivirus were performed as in previous reports^19, 20^. Briefly, the pseudo-typed lentiviruses were generated by co-transfecting 293T cells with the constructed plasmids and two packaging vectors (pCMV Delta R8.2 and pCMV-VSVG). The packaged lentivirus particles were purified using 8% PEG-8000 and 0.5 M NaCl. And they were used to infect MSCs or MNCs, and screened by puromycin. The generated stable cells were confirmed by qRT-PCR examination.

### Bone resorption pit assays

The bone slices were made from ox bone and placed into the 24-well plate according to the previous study^17^. Primary MNCs were plated in the plates (Corning, USA) and incubated with the osteoclast-induced medium. At day 8, the plate was washed with PBS to remove the cells, and washed three times with distilled water. The bone slice was incubated with 1% toluidine blue in 1% sodium borate droplet. The sizes of the pit were evaluated under a microscope, and the data was analyzed using Image J software.

### RNA immunoprecipitation (RIP)

RNA immunoprecipitation (RIP) was performed according to our previous study^21^. Briefly, beads were incubated with 5µl antibody against EZH2 or IgG at 4 [overnight. After washing twice with RIPA lysis buffer, the beads were transferred to tubes with cell lysates and then rotated at 4 [for 6h. RNA extraction from the beads was further collected using TRIzol reagent and analyzed by qRT-PCR examination. The results were normalized with the input control.

### Co-Immunoprecipitation (Co-IP)

The cells were lysed by cold lysis buffer, and the lysate was collected and centrifuged. The supernatant was qualified by the BCA assay kit (Thermo Fisher Scientific) and incubated with indicated antibodies at 4 [for 4h. Then, the protein A+G agarose was added to the lysate at 4[. After 1h incubation, the agarose were washed with cold washing buffer for three times, and protein loading buffer was added. The mixture was boiled and subjected to western blotting analyses.

### ChIP assay

Chromatin was extracted from pVector and pHottip infected MSCs and fast chromatin immunoprecipitation (ChIP) was performed using Protein A+G agarose (Beyotime, Shanghai)^22^. The following antibodies were used for ChIP: H3K9me3 (1:100; Cell Signaling Technology, USA) and H3K27me3 (1:100; Cell Signaling Technology, USA). The nonimmune IgG control samples were used as a mock. ChIP-qPCR primers are listed in Supplementary Table 2. Fold enrichment was calculated using the following equation based on Ct as 2^ΔCt^, where ΔCt = (Ct^mock^ – Ct^Target^).

### Mouse femoral fracture model and Hottip modified MSCs treatment

To assess the bone fracture healing capacity of Hottip, age-matched WT and KO mice (n=8) were utilized in the fracture model. A mid-shaft transverse osteotomy was performed using an electric saw, followed by fixation with a 0.014-inch Kirschner’s needle inserted into the intramedullary canal under general anesthesia^23^. The incision was closed, and the fracture was confirmed by X-radiography. For investigating the therapeutic effect of pHottip-infected MSCs on bone fracture, 16 C57BL/6J mice were randomly assigned to two groups post-surgery: MSCpVector and MSCpHottip. Modified MSCs (10^7^ cells/ml) were injected into the fracture sites^24^, and the animals were returned to standard husbandry. X-ray radiography was performed at week 2 and week 4 using a digital X-ray machine (Faxitron X-Ray Corp., USA) to evaluate the fracture healing effect. Animals were randomly selected for sacrifice, and their femurs were harvested for further analysis.

### Micro-computed tomography (μCT)

The microstructure of mouse femur, tibia and centrum, as well as the callus on the fracture sites were quantitatively assessed by μCT scanning as previously described^25^. Reconstruction was carried out using different thresholds (low attenuation=158, high attenuation=211) with SkyScan™ NRecon (Bruker, Skyscan1172, Belgium) software^26^. Bone volume (BV), tissue volume (TV), bone volume/total volume (BV/TV), bone surface/bone volume (BS/TV) and trabecular thickness (Tb.Th) were recorded for quantitative analyses.

### Statistics

Statistical analyses were performed using IBM SPSS statistics 24. The two-tailed Student’s t-test was used to determine the significance between two groups. Two-way ANOVA followed by Tukey’s test was performed to compare the data among multiple groups. All data are presented as the mean ± SD of at least three independent experiments. P < 0.05 was used to indicate statistical significance.

## Results

### Abnormal bone metabolism was found in Hottip KO mice

To investigate the role of Hottip in the musculoskeletal system, we conducted global metabolomic profiling between WT and KO mice using LC-MS. A total of 50 and 139 metabolites were identified in negative and positive ion modes, respectively (Figure 1a and Supplementary Figure1a). Heat map analysis revealed differentially expressed metabolites in both modes (Figure 1b and Supplementary Figure1b). Notably, cystine, L-cysteine-glutathione disulfide, cytidine 5’-diphosphocholine, LysoPE 18:2, PE (2:0/18:1), and PS (18:0/18:1) were prominently altered, suggesting dysregulation of histidine metabolism, steroid hormone biosynthesis, and beta-alanine metabolism in KO mice (Figure 1c).

**Figure 1.**
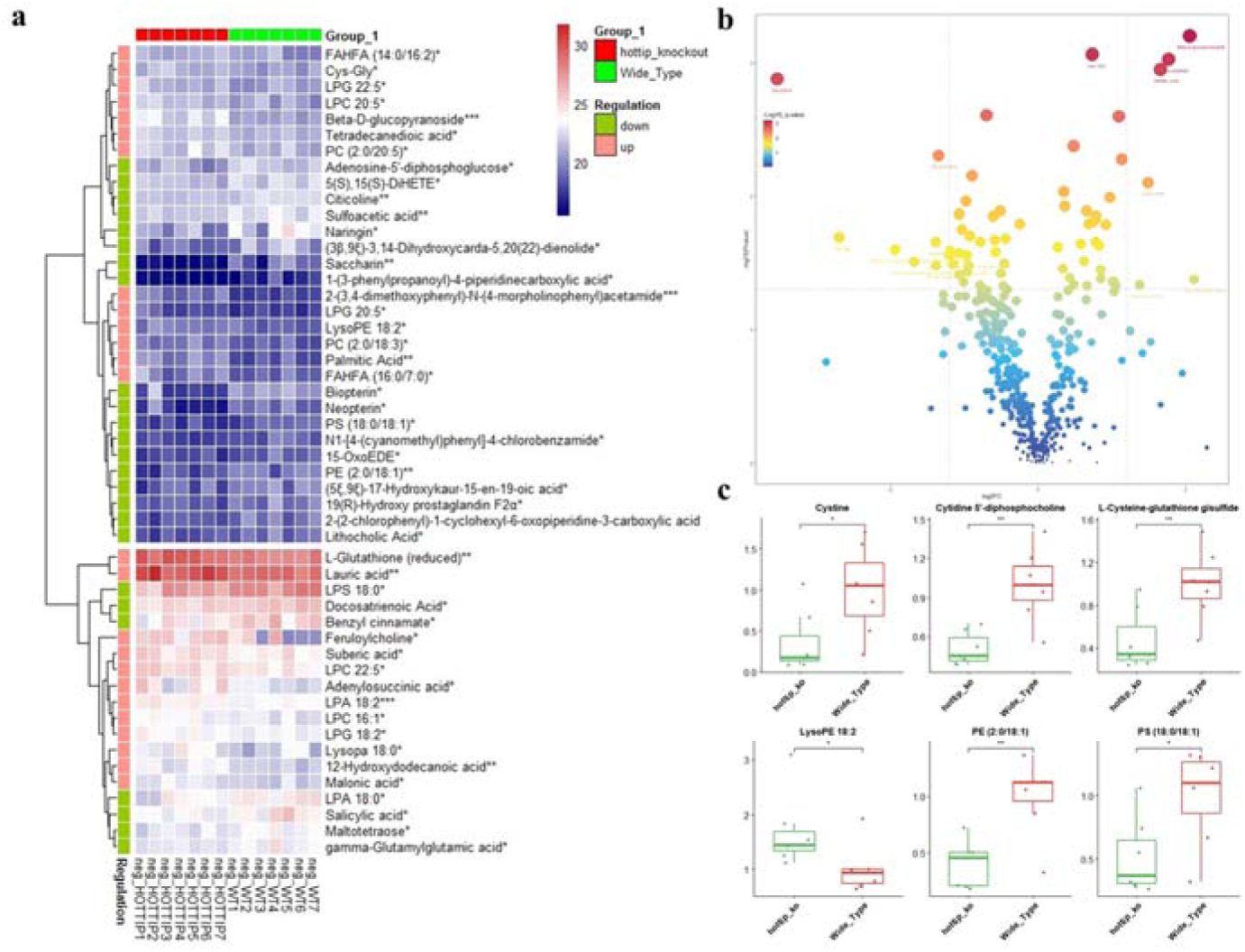
Bone metabolomic profiling was analyzed between WT and Hottip KO mice. (a), Heatmap showed the differently expressed metabolites between WT and Hottip KO mice using liquid chromatography mass spectrometry. (b), The volcano plot of all the different metabolites. P<0.05 was set as the cut-off criterion of a significant difference, and the color and shape of the dots were associated with -log10 value. (c), the representative metabolites associated with skeletal homeostasis. Data are presented as the mean±SEM from seven independent experiments. n=7; *, P<0.05; **, P<0.01; ***, P<0.001.

Furthermore, we compared the metabolites between KO mice and postmenopausal osteoporotic mice^26^. Enrichment analysis of differential metabolites indicated consistent dysregulation of alanine metabolism, fatty acid biosynthesis, and glutathione metabolism in both groups (Supplementary Figure 1c). These findings suggest that Hottip may play a role in bone regeneration and development by influencing bone metabolism.

### Abnormal bone tissues and impaired bone quality were observed in Hottip KO mice

To identify the actual contribution of Hottip in bone regeneration, we compared the bone tissues derived from Hottip KO mice and WT groups. By micro-CT examination, the thinner cortical and looser cancellous bone were shown in the representative 3D images of femur, tibia (Figure 2a) as well as centrum (Supplementary Figure 2a) derived from KO mice. The quantitative assays revealed a significant decrease in BV/TV, BS/TV and Tb.Th in KO mice (Figure 2b-2c, Supplementary Figure 2b). The further H&E staining showed less trabecular bone mass around the metaphyseal region KO mice (Figure 2d and 2f, Supplementary Figure 2c). Moreover, TRAP staining indicated more TRAP positive osteoclasts in femurs of KO mice (Figure 2e). We also observed the similar phenomena in the tibia and centrum derived from KO mice (Figure 2f, Supplementary Figure 2d). Based on the IHC staining, the decreased OCN and OSX expression were also found in Hottip KO mice (Figure 2g-2h). These results suggest that Hottip contributes to bone regeneration, possibly *via* mediating bone formation and resorption.

**Figure 2.**
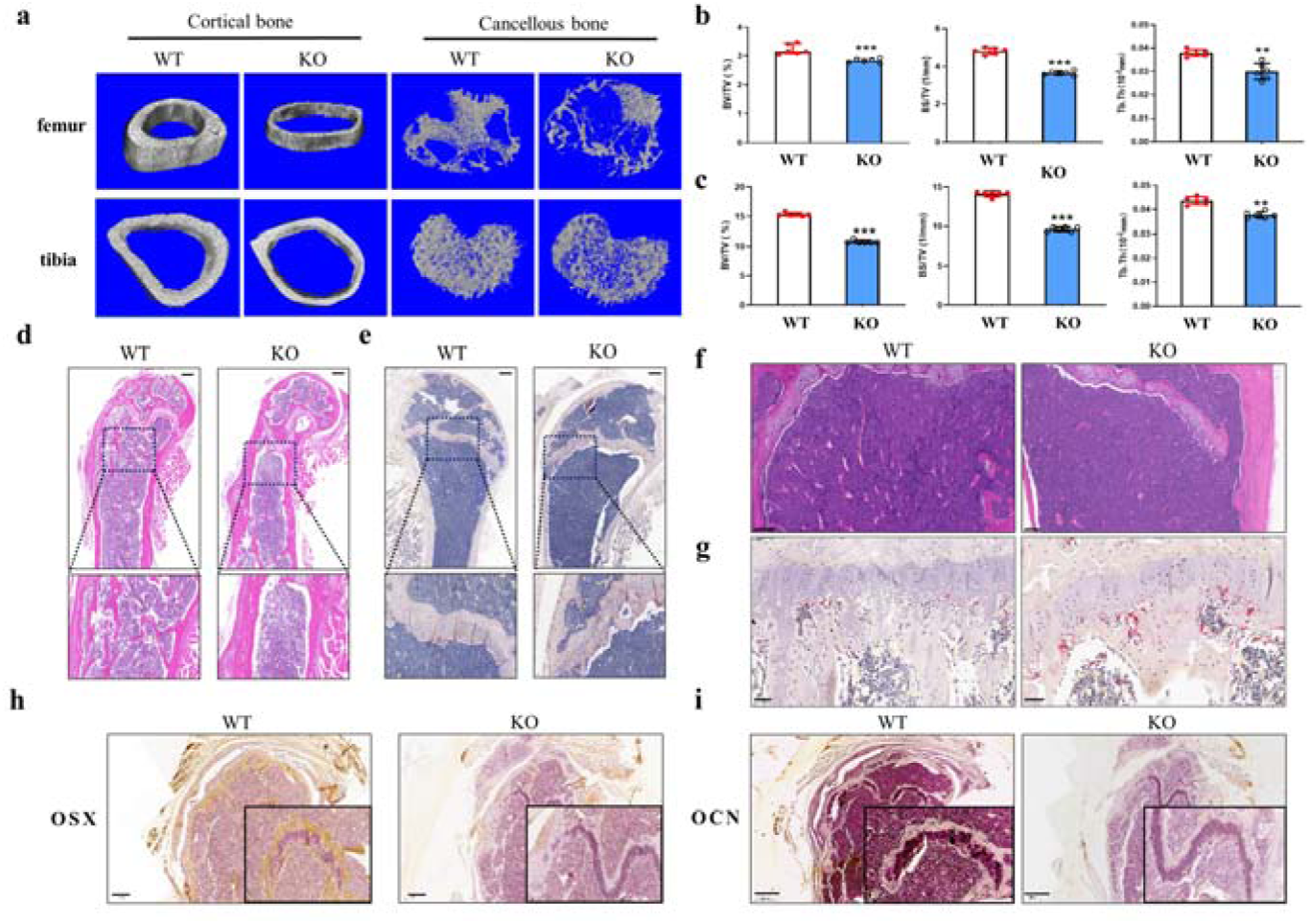
Abnormal bone microarchitecture and impaired bone quality was found in Hottip KO mice. (a), The cortical and cancellous microarchitecture of femurs and tibia of Hottip KO mice were examined by µCT and the representative 3D images were reconstructed. (b-c), Quantitative analyses of bone volume over total volume (BV/TV), bone surface over total volume (BS/TV) and Tb thickness (Tb. Th) in cancellous femurs (b) and tibias (c). (d-f), H&E staining and TRAP staining of sagittal plane of the metaphyseal region of femurs and tibias. (g-h), The expression of Osx (g) and OCN (h) in femurs derived from KO mice was examined by immunohistochemistry staining. **, P<0.01; ***, P<0.001.

To investigate the role of Hottip in bone regeneration, we compared bone tissues from Hottip knockout (KO) mice with wild-type (WT) groups. Micro-CT examination revealed thinner cortical and looser cancellous bone in KO mice, along with significant decreases in BV/TV, BS/TV, and Tb.Th. H&E and TRAP staining showed reduced trabecular bone mass and more TRAP-positive osteoclasts in KO mice. Additionally, IHC staining indicated decreased expression of OCN and OSX in Hottip KO mice. These findings suggest that Hottip plays a crucial role in bone regeneration, potentially by mediating bone formation and resorption.

### A delayed fracture healing was found in Hottip KO mice

To further confirm the *in vivo* role of Hottip, we established a mouse femoral fracture model to compare the healing process between KO and WT mice. The fracture healing was monitored by X-rays examination, and the results showed that the gaps between the fracture sites almost disappeared in WT mice while it remained visible in KO mice at week 2 and week 4 (Figure 3a). We also found that the newly formed callus of WT group was thicker at week 2 and the callus was absorbed and the gap disappeared at week 4 when compared with KO group (Figure 3a), indicating that the fracture healing process was delayed in this KO mice. The calluses were assessed by μCT scanning and 3D reconstructed images also confirmed the thinner newly formed callus in KO mice at week 4 (Figure 3b); and the quantitative examination showed that the KO group had significantly decreased BV/TV and BS/TV, suggesting less newly formed mineralized bone in this group (Figure 3c). The H&E and Masson staining of fracture sites revealed that less osteoblasts were converged and the bone regeneration was less vigorously in KO mice when compared with WT group (Figure 3d). Furthermore, the decreased OCN and OSX expression were also observed in KO group by IHC staining (Figure 3d), suggesting an impaired effect of Hottip knockout on bone formation and regeneration. Moreover, the three-point bending mechanical testing was performed and results showed that the Hottip KO group had a significant decrease in ultimate load, E-modulus and Yield strength (Figure 3e).

**Figure 3.**
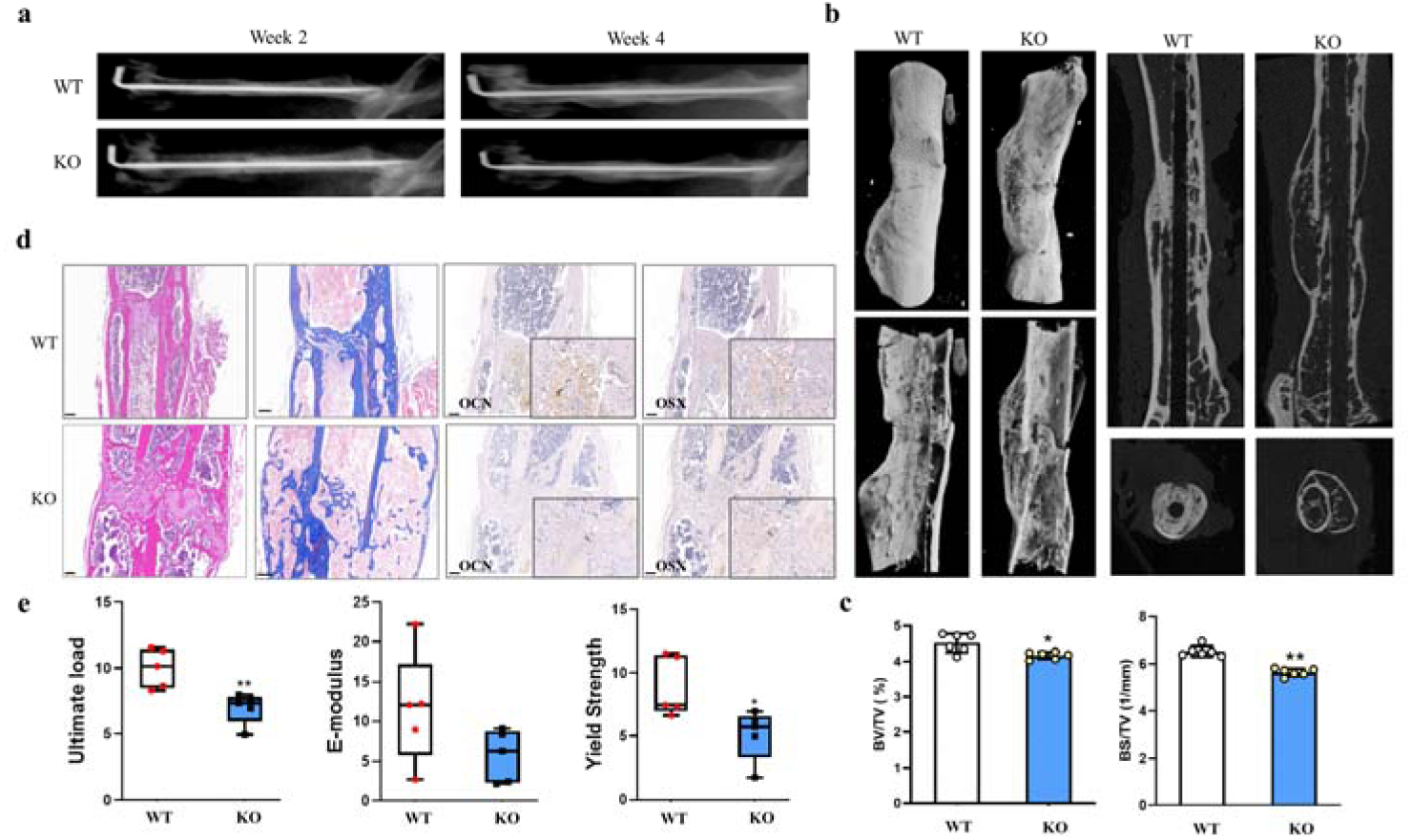
The delayed fracture healing was observed in Hottip KO group. (a), The fractures were monitored by X-rays examination at week 2 and week 4. (b), Three-dimension μCT images of the mouse femur fracture zone were taken at week 4 after surgery. (c), Quantitative analyses of bone total volume and over total volume (BV/TV) and bone surface/bone volume (BS/TV). (d), H&E, Masson staining and IHC staining for OSX and OCN. (e), E-modulus, ultimate load and yield strength were calculated though three-point bending mechanical testing. *, P<0.05; **, P<0.01.

### Hottip maintained skeletal homeostasis by promoting osteogenic differentiation and inhibiting osteoclastogenesis

Firstly, we investigated the osteogenic potential of MSCs derived from KO mice. During osteogenic differentiation, the early marker alkaline phosphatase (ALP) activity was obviously decreased at day 7 in KO-MSCs (Figure 4a), and fewer mineralized nodules was also observed at day 21 (Figure 4b). Furthermore, the expression of osteogenic marker genes including Runt-related transcription factor 2 (Runx2), ALP, bone morphogenetic protein 2 (BMP2), osteocalcin (OCN), osteopontin (OPN) and osterix (Osx) were significantly repressed in the KO group (Figure 4c). To further confirm the function of Hottip in osteogenesis, the lentiviral system was used to stably reinforce or silence Hottip expression in mouse MSCs, and generated the stable cells, which were verified by qRT-PCR examination (Supplementary Figure 3a-3b). Consistently, the results of ALP activity, calcium nodules and marker gene expression revealed that Hottip knockdown significantly suppressed osteogenic differentiation of MSCs (Figure 4d-4f). On the contrary, the ectopic overexpression of Hottip sharply stimulated ALP activity (Figure 4g), improved mineralized nodules (Figure 4h) and promoted the crucial marker genes expression (Figure 4i). Rescue assays were conducted to clarify the essential role of EZH2 in Hottip-mediated osteogenic differentiation. In vivo experiments revealed that overexpression of EZH2 partially counteracted the bone-regenerative effects of Hottip (Supplementary Figure 4a-4d). Furthermore, EZH2 was found to be vital in mediating the transcription of osteogenic genes as evidenced by partial inhibition of osteogenic marker genes expression following overexpression of Hottip. (Supplementary Figure 4e). Collectively, all these data suggest that Hottip may facilitate osteogenic differentiation of MSCs, overexpression of EZH2 could partially abolished Hottip-promoted bone regeneration.

**Figure 4.**
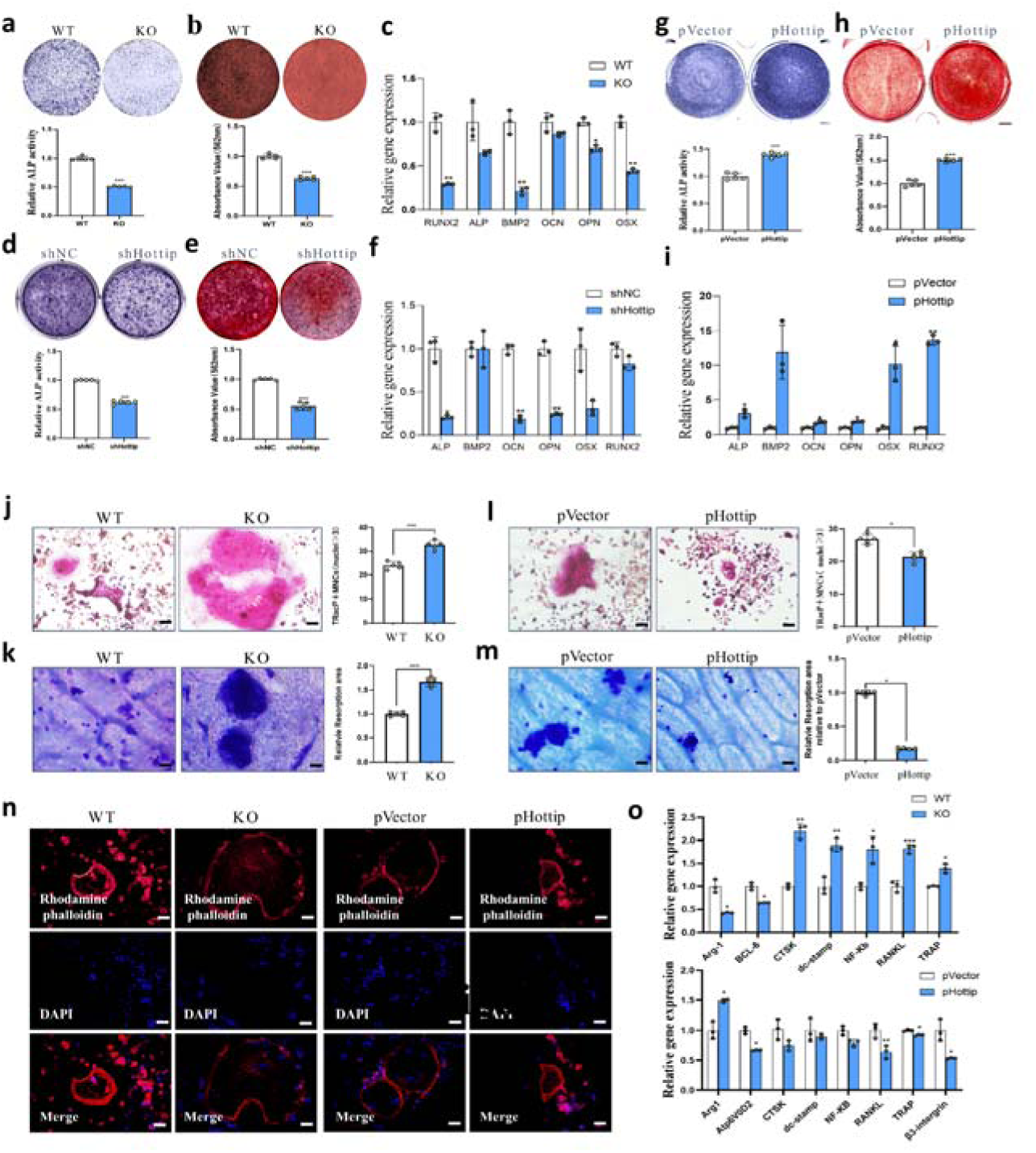
Hottip maintained skeletal homeostasis by promoting osteogenic differentiation and inhibiting osteoclastogenesis. (a-c), Under osteogenic induction, ALP activity (a), calcified nodules (b) and osteogenic marker genes expression (c) were all suppressed in MSCs derived from Hottip KO mice. (d-f), ALP activity (d), calcified nodules formation (e) and osteogenic marker genes expression (f) were all decreased by shHottip. (g-i), ALP activity (g), calcified nodules formation (h) and osteogenesis marker genes expression (i) were promoted by pHottip in MSCs. n=3, (j), Qualitative and quantitative examination of TRAP positive osteoclast (≥3 nuclei) in Hottip KO and WT mice. (k), Qualitative and quantitative examination of bone resorption. (l), The staining of rhodamine phalloidin for F-actin ring in osteoclasts. (m), Qualitative and quantitative examination of TRAP positive osteoclast (≥3nuclei) in pHottip and pVector groups. (n), Qualitative and quantitative examination of bone resorption in the Hottip overexpressing cells. (o), The expression of several markers related to osteoclast differentiation was measured by qRT-PCR assays at day7. *, P<0.05; **, P<0.01; ***, P<0.001.

On the other hand, the osteoclastogenic effect of Hottip was examined and it was found the bigger and mature osteoclasts in MNCs derived from Hottip KO mice by TRAP staining (Figure 4j). Their bone resorption was strongly promoted as well (Figure 4k). Conversely, Hottip overexpression suppressed osteoclastogenic differentiation and bone resorption (Figure 4l-4m, Supplementary Figure 3c). Considering that F-actin ring formation plays a critical role in osteoclastic bone resorption, we investigated the influence of Hottip on F-actin ring formation. As shown in Figure 4n, Hottip knockout stimulated while its overexpression inhibited F-actin ring formation in osteoclasts. Furthermore, several crucial genes related to osteoclast differentiation such as CTSK, TRAP, β3-intergrin, Atp6v0d2, Arg-1, BCL-6, dc-stamp, NF-ΚB and RANKL were examined and the results showed that they were significantly promoted by Hottip knockout while obviously suppressed by Hottip overexpression (Figure 4o). Taken together, these results suggest that Hottip possibly serves as an negative regulator of osteoclast differentiation and bone resorption.

### Hottip suppressed EZH2 expression and histone methylatic modification

To better understand how lncRNA Hottip regulates bone homeostasis, the transcriptome alteration between KO and WT mice was conducted. Bioinformatics analysis of the RNA-seq data indicated the differently expressed gene mainly focused on developing skeletal muscle tissue and organ (Figure 5a). The further gene set enrichment analyses revealed that Polycomb Repressive Complex 2 (PRC2) members were conspicuously influenced by HOTTIP knockout (Figure 5b). The expression of PRC2 members, including EED, SUZ12 and EZH2, were confirmed by qRT-PCR examination, and EZH2 was the most changeable (Figure5c). Consistently, Hottip overexpression suppressed while its knockdown promoted EZH2 expression in MSCs and MNCs at the protein level (Figure 5e-5g, Supplementary Figure 5a-5d). The further IHC staining also revealed the decreased EZH2 expression in the femur and tibia derived from KO mice (Figure 5d). Considering that EZH2 is a highly conserved histone methyltransferase that specifically catalyzes histone H3K9 and H3K27 for trimethylation^27, 28^, we thereby examined the expression of H3K27me3 and H3K9me3 in MSCs or MNCs. It was found that Hottip knockdown stimulated while Hottip overexpression suppressed the expression of H3K27me3 and H3K9me3 in MSCs (Figure 5e). As expected, the promoted expression of H3K27me3 and H3K9me3 was observed in the MNCs derived from KO mice (Figure 5f), while the suppressed expression of H3K27me3 and H3K9me3 was found in Hottip overexpressing MNCs with or without RANKL (Figure 5g). All these data suggested that Hottip mediated histone trimethylation *via* suppressing EZH2 expression.

**Figure 5.**
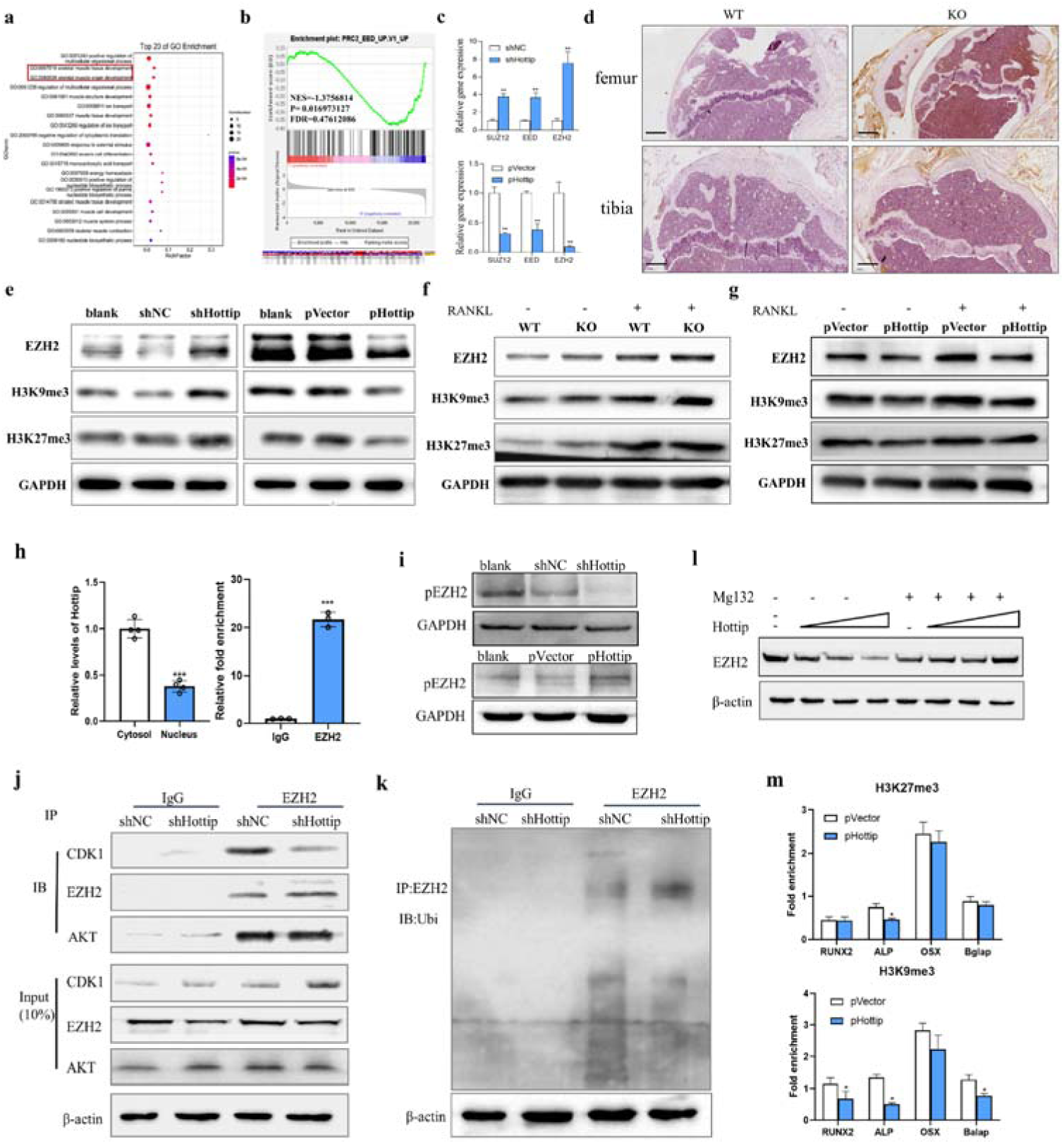
Hottip mediated bone homeostasis by *recruiting EZH2 and mediating its stability*. (a), RNA-seq data indicated that the differently expressed gene mainly focused on developing skeletal muscle tissue and organ. (b), GSEA revealed that Polycomb Repressive Complex 2 (PRC2) members were the most remarkable genes. (c), qRT-PCR examination was applied to validate the expression alteration from RNA-seq data. (d), The protein level of EZH2 was monitored by IHC staining in femur and tibia specimens. (e), The protein level of EZH2, H3K9me3 and H3K27me3 was examined by Western blot in shHottip and pHottip infected MSCs. (f-g), The protein levels of EZH2, H3K9me3 and H3K27me3 were examined by Western blotting in shHottip and pHottip infected BMMs with or without RANKL. (h), In order to investigate the subcellular distribution of Hottip and its interaction with EZH2, we utilized the FastPure Cytoplasmic & Nuclear RNA Purification Kit and performed RNA immunoprecipitation assays, respectively. (n=3). (i), Western blot analysis of phosphorylated EZH2 proteins in Hottip knockdown and overexpressing MSCs (n=3). (j), The protein levels of endogenous EZH2-associated CDK1 and Akt in control and Hottip silenced cells were determined by co-immunoprecipitation assay (n=3). (k), Western blot of endogenous EZH2-associated ubiquitination in control and Hottip silenced cells after immunoprecipitation with anti-EZH2 antibody in MSCs (n=3). (l), The EZH2 protein level was detected by western blot in MSCs after treatment with MG132 (10μM) for 6h (n=3). (m), ChIP assays showed the relative enrichment of H3K27me3 and H3K9me3 on the promoters of several osteoblast and osteoclast gene such as RUNX2, ALP, OSX and Balap promoter after overexpression of Hottip (n=3). *, P<0.05; ***, P<0.001.

### Hottip recruited EZH2 and mediated EZH2 degradation

It has demonstrated that EZH2 physically interacted with several lncRNAs and triggered gene silence, and we therefore wondered whether Hottip might directly associate with EZH2 and thus mediated its stability. To testify this hypothesis, we performed RNA immunoprecipitation (RIP) assay and found that Hottip was preferentially enriched in the EZH2-recruited complex (Figure 5h). As many lncRNAs have been reported to modulate the phosphorylation and stability of their binding proteins, we also examined whether Hottip could mediate the phosphorylation of EZH2. As expected, the expression of phosphorylated EZH2 was enhanced by Hottip overexpression whereas it was suppressed by Hottip knockdown (Figure 5i). In terms of that CDK1 and Akt are two major kinases that mediate the phosphorylation of EZH2 and suppress its methyltransferase activity, we next examined the CDK1-EZH2 and Akt-EZH2 interaction by co-IP. We then immunoprecipitated endogenous EZH2 protein from Hottip silenced MSCs and control cells to examine the CDK1-EZH2 and Akt-EZH2 interaction. It was found that EZH2 pulled down less CDK1 protein in the Hottip knockdown cells when compared to that in control cells (Figure 5j). However, the silence of Hottip did not affect the mutual interaction between Akt and EZH2 (Figure 5j). Considering that CDK1 mediated phosphorylation of EZH2 and then stimulated its ubiquitination and subsequent degradation^29^, we utilized EZH2 antibody to pull down endogenous EZH2 proteins and examined their ubiquitin modification. The co-IP results revealed an increased EZH2 ubiquitination in shHottip infected cells (Figure 5k). As well known, the ubiquitin-proteasome pathway tightly modulates the protein stability. The proteasome inhibitor MG132 was used to investigate the rescued effect of Hottip overexpression on EZH2 expression. Immunoblotting results showed that the enhanced expression of Hottip decreased endogenous EZH2 in a dose-dependent manner, whereas this decrease was alleviated by MG132 treatment (Figure 5l, Supplementary Figure 6). This result proposed that destabilization of EZH2 mediated by Hottip may depend on the proteasomal degradation. We therefore concluded that Hottip recruited EZH2 and induced EZH2 phosphorylation and degradation, which led to suppressing histone methylation modification. Given that EZH2 epigenetically suppresses gene expression through specifically inducing methylation on Lys-9 and Lys-27 of histone 3 (H3K9me3 and H3K27me3), we conducted chromatin immunoprecipitation (ChIP) assays to examine whether Hottip mediated the interaction between H3K9me3 as well as H3K27me3 and osteogenic genes’ promoter. According to qRT-PCR results, Hottip overexpression significantly suppressed the DNA fragment from RUNX2, ALP and Balap promoter enriched by H3K9me3. Also, it suppressed the DNA fragment from ALP promoter enriched by H3K27me3 as well (Figure 5m), suggesting Hottip could inhibit the binding ability of H3K9me3 and H3K27me3 with these osteogenic genes’ promoter and stimulate bone formation.

To further clarify the essential role of EZH2 in Hottip-mediated osteoblast differentiation, a rescue study was conducted and the results showed that overexpression of EZH2 partially counteracted the Hottip-promoted osteogenic effects (Supplementary Figure 4a-4d) as well as the Hottip-induced upregulation of the osteogenic genes (Supplementary Figure 4e).

### Hottip modified MSCs accelerated bone fracture healing *in vivo*

To further identify the clinical significance of Hottip, a mouse femoral fracture model was established and the Hottip overpressing mesenchymal stem cells (MSC^pHottip^) were locally injected into the fracture sites. X-ray images were taken at week 2 and week 4, and the results showed that the gap in the fracture sites nearly disappeared at week 4 in the MSC^pHottip^ group (Figure 6a). Further micro-CT scanning showed more newly mineralized calluses in this treated group (Figure 6b). The quantification of BV/TV and BS/TV were analyzed and they were significantly increased in this overexpressing group (Figure 6c), indicating more newly formed bone with MSC^pHottip^ treatment. Further HE, Masson and IHC staining were performed to evaluate the histological properties of newly mineralized-bone tissues. As shown in Figure 6d and 6e, the regeneration of fracture sits was more quickly and vigorously in MSC^pHottip^ group when compared with the control group at week 4 post-surgery. Moreover, increased OSX and OCN expression were also observed in MSC^pHottip^ treated group (Figure 6f). All these data indicated that Hottip promoted bone regeneration and facilitated fracture healing. To sum up, a regulatory axis was proposed to illustrate the underlying molecular mechanism (Figure 6g): Hottip physically interacted with EZH2, and subsequently potentiated CDK1-EZH2 interaction, leading to the phosphorylation and ubiquitination of EZH2. The Hottip-mediated EZH2 degradation facilitated the activation of histone methylation modification, which eventually promoted bone formation and inhibited bone resorption.

**Figure 6.**
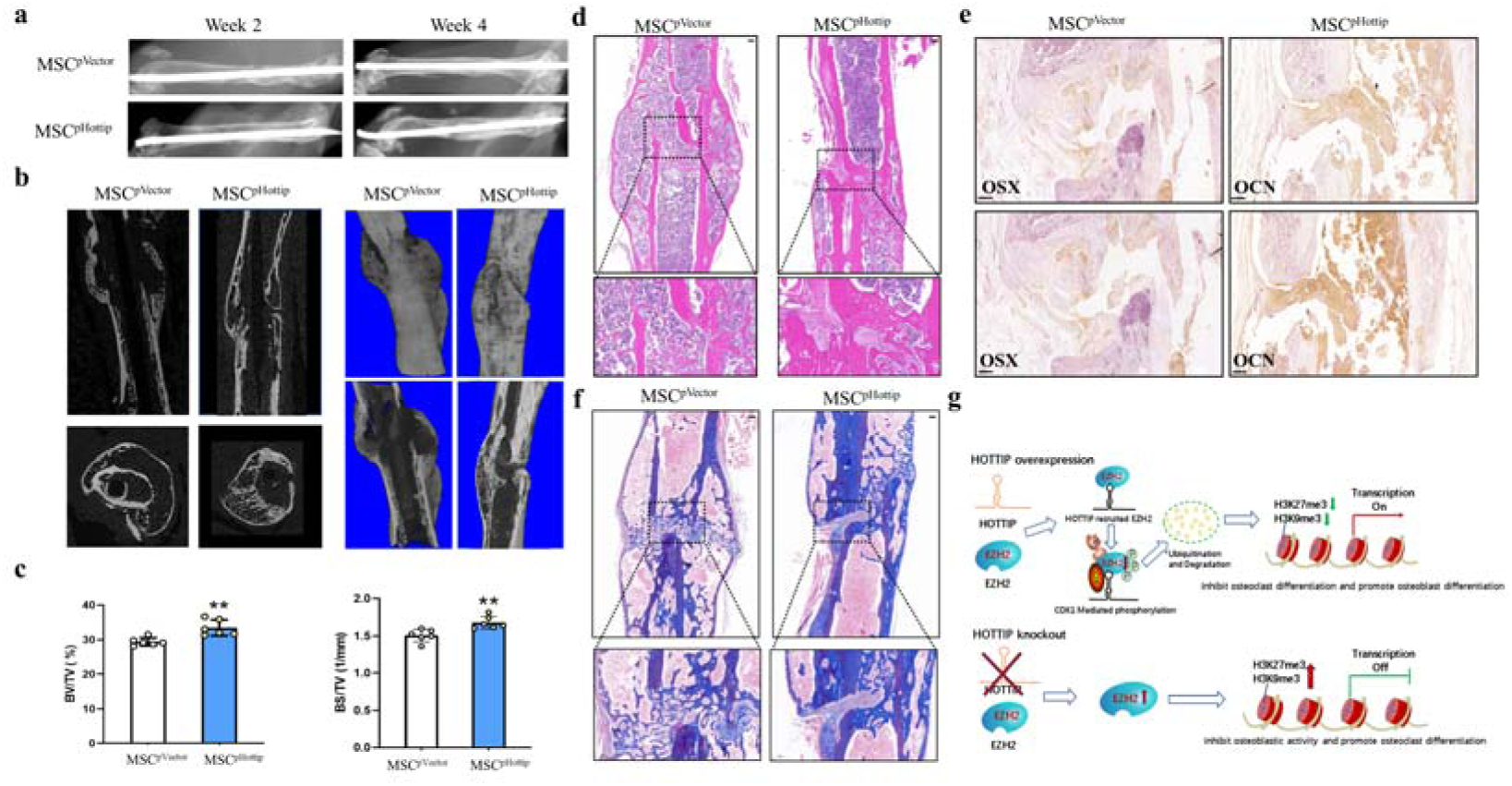
pHottip modified MSCs accelerated bone fracture healing in the mouse. (a), X-ray was taken at week 2 and week 4, and representative images showed that the bone remodeling was more vigorous in pHottip group. (b), Three-dimension μCT images confirmed better fracture healing condition and more continuous callus in pHottip group. (c), The statistical diagram of BV/TV and BS/TV were calculated in the two groups. (d), The H&E staining showed better repaired effect in pHottip group. (e), The Masson staining showed better repaired effect in pHottip group. (f), the expression of Osx and OCN were examined by IHC staining in the new bone zone in pHottip group. (g), The hypothetical model of Hottip regulatory network. **, P<0.01

## Discussion

Bone remodeling is essential for bone homeostasis, which depends on the resorption of bones by osteoclasts and the formation of bones by the osteoblasts. The imbalance of this tightly coupled process causes abnormal bone regeneration, resulting in bone related diseases such as osteoporosis. Thus, unraveling the underlying mechanism of interaction between osteoclasts and osteoblasts is critical for bone biology. In recent years, lncRNAs’ emergence provides novel insights for this interplay. Although lncRNAs have been demonstrated to be widely involved in bone homeostasis^30–32^, few lncRNAs were reported to serve as a mediator to balance the osteoblast/osteoclast differentiation. In the present study, we firstly identified lncRNA Hottip as a switch to maintain the balance of bone homeostasis *via* promoting osteoblast differentiation and suppressing osteoclast differentiation.

HOTTIP originates from a genomic region in the 5’tip of the HOXA locus^33^. Its genomic location endowed it with a role in activating numerous 5’HOXA genes. Accumulating evidence demonstrated that HOX genes were critical for tissue formation and regeneration^34, 35^, and their aberrant activation or inhibition may affect bone homeostasis. We therefore speculated that Hottip played a similar role in bone homeostasis as its host gene. To validate this hypothesis, comprehensive analyses of metabolomics and transcriptomics were conducted between Hottip KO and WT mice. We found that the differently-expressed metabolites were mainly enriched on the beta-alanine metabolism, fatty acid biosythesis, alanine aspartate, glutamate metabolism and glutathione metabolism, which were consistent with the osteoporotic mice. These data suggest that these KO mice may exhibit an osteoporotic phenotype or be prone to osteoporosis. On the other hand, amino acid metabolites, especially, cystine and L-cysteine were obviously decreased in KO mice, which subsequently inhibited insulin secretion and reduced the effects of insulin-like growth factor 1 (IGF-1) on osteoblast differentiaiton^36^. And the phosphatidylethanolamine was sharply reduced in KO mice, and this metabolite was reported to induce bone formation and improve cell metabolism and osteogenic differentiation^37^. As for the transcriptomics examination, the bioinformatics analysis showed that Hottip knockout affected several signaling pathways, and osteoclast differentiation was the most striking one. The further bone microstructure and histomorphometry examination validated that osteoclasts were strongly activated in the KO mice. The *in vitro* cell data also confirmed that Hottip silence promoted osteoclast differentiation. What’s more, we also demonstrated that Hottip KO impaired bone formation and suppressed osteoblast differentiation. Based on these data, Hottip was considered as a switch to maintain the interaction between bone formation and resorption.

Recently, it is recognized that lncRNAs can recruit protein complexes and modulate protein function, which conversely determines lncRNA functional effects^38^. To identify the protein targets, the RNA-seq analysis revealed that Polycomb Repressive Complex 2 (PRC2) members were potential targets of Hottip. The previous study have demonstrated that lncRNAs might exert their function through physically interacting with Polycomb Repressive Complex 1 (PRC1), Polycomb Repressive Complex 2 (PRC2) or other RNA binding proteins, which provided strong support for our hypothesis. The bioinformatics prediction and RIP experiment confirmed that the PRC2 member EZH2 could be recruited by Hottip. We also found this interaction improved the mutual interaction between EZH2 and Akt, which induced EZH2 ubiquitination and degradation, thereby leading to suppression of EZH2.

As one of the most significant epigenetic modifications, histone lysine methylation has emerged as a critical player in regulating gene expression and their stability. Increasing evidence demonstrated that histone methylation is closely associated with bone remodeling and regeneration. Many lncRNAs have been demonstrated to participate in this epigenetic modification^39^. LncRNA Hottip was reported to interact with the WDR5 protein directly and recruit the WDR5/MLL complexes across the HOXA locus, driving H3K4 methylation and the transcription of HOXA gene^40^. This interaction also promoted the translocation of WDR5 into the nucleus and stimulated the transcription of β-catenin^41–42^ As a core component of PRC2, EZH2 epigenetically silences its target gene transcription by specially catalyzing the trimethylation of histone H3 lysine 27 (H3K27me3) and histone H3 lysine 9 (H3K9me3). The EZH2/histone methylation axis is well-recognized signaling to mediate the bone formation and resorption. Silence of EZH2 has been reported to stimulate the key components of osteogenic signaling, such as BMP2 pathway cascades. Its inhibitor GSK126 enhanced osteoblast differentiation by disrupting the H3K27 methylation^43^. More importantly, EZH2 activity and H3K27me3 levels were upregulated in the RANKL-induced pre-osteoclasts, and blocking EZH2 methyltransferase activity could suppress the formation of mature osteoclasts^44^. EZH2 was thereby considered as an inhibitor of osteoclastogenesis and activator of osteoclastogenesis^45^, indicating it can maintain the balance between bone formation and resorption. Our results demonstrated that Hottip overexpression suppressed while its silence promoted the expression of EZH2 as well as H3K9me3 and H3K27me3 in osteoblast and osteoclast, which was consistent with the regulatory effects of Hottip on bone formation and resorption. To further address the role of Hottip in the histone 3 methylation mediated osteogenesis, a ChIP assay was conducted and we found that Hottip decreased the enrichment of H3K9me3 and H3K27me3 at the RUNX2, ALP, OSX and Balap promoter in MSCs, which led to the stimulation of bone formation.

Collectively, our study depicted a novel regulatory role of lncRNA Hottip in bone regeneration. It was found that Hottip promoted osteoblast differentiation and suppressed osteoclast differentiation, indicating it may serve as a switch to maintain the bone homeostasis. Further mechanism investigation revealed the Hottip physically interacted with EZH2 and mediated its stability, eventually leading to the modification of histone methylation. This finding might provide a bright insight in developing Hottip as a potential therapeutic target for bone regeneration.

**Table 1.**
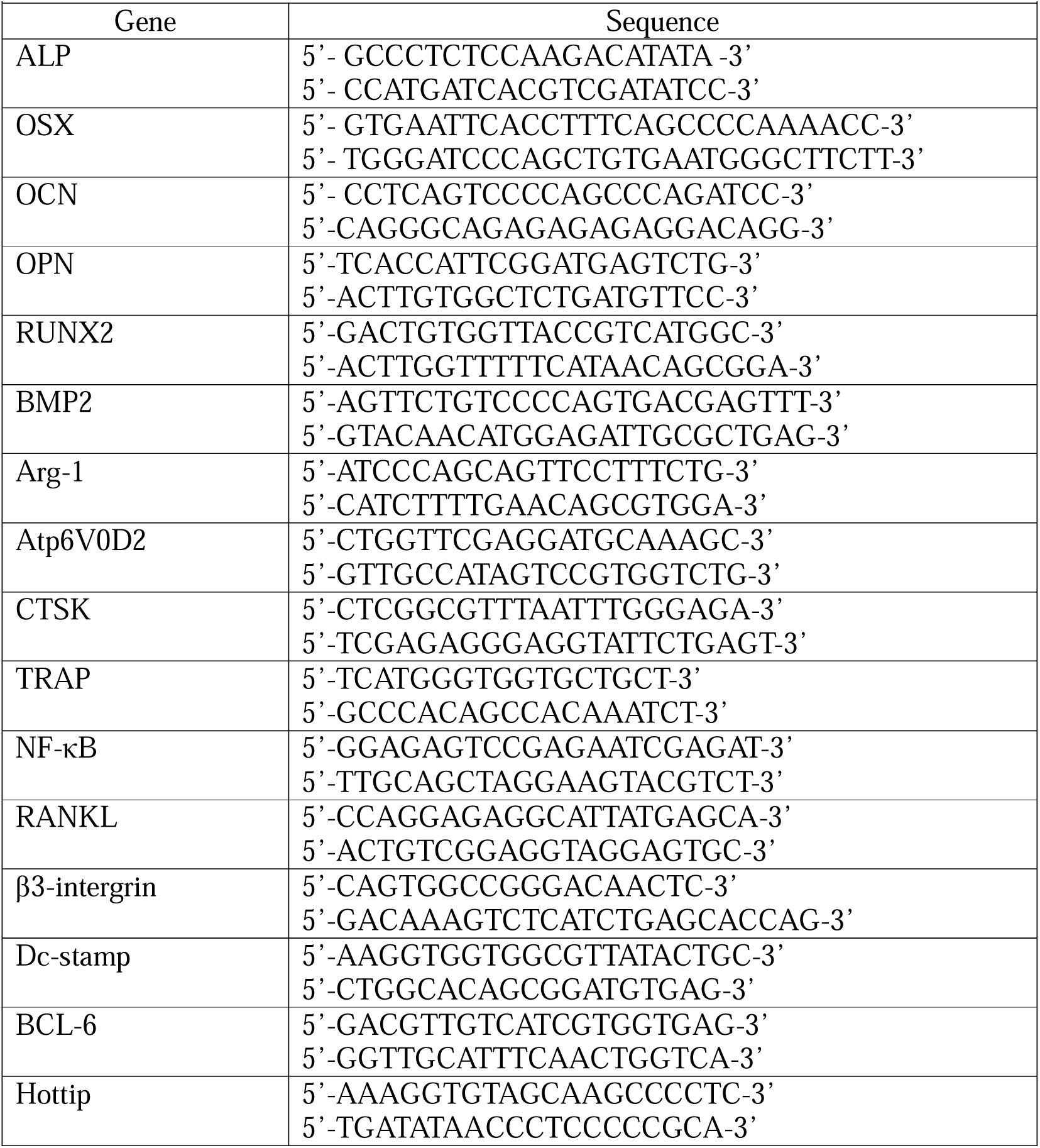
Primer sequences for qPCR.

**Table 2.**
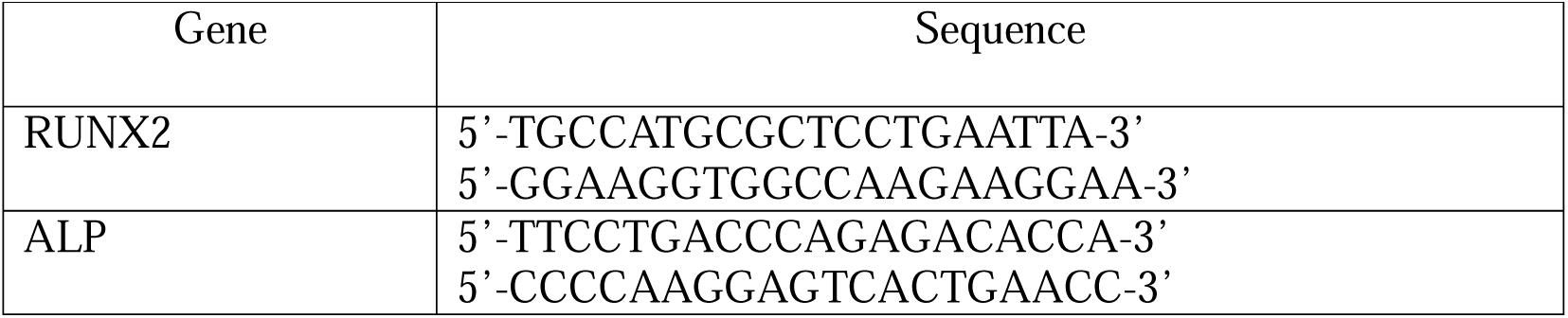
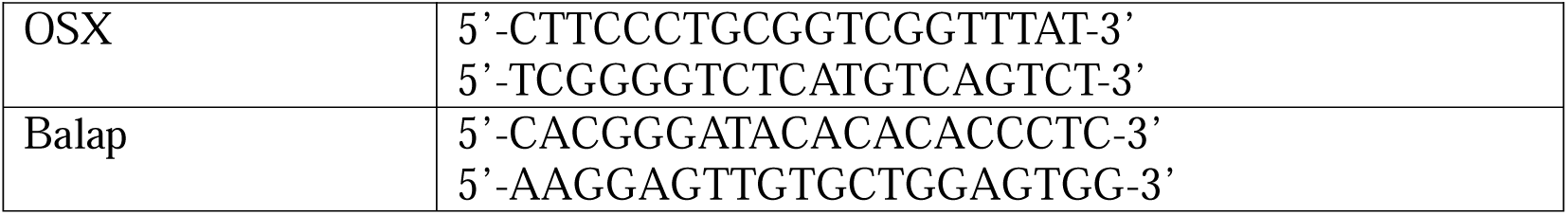
Primer sequences for CHIP-PCR.

## Funding supports

This work was supported by grants from the National Natural Science Foundation of China (No. 81772404).

---

**Supplementary Figure 1.**
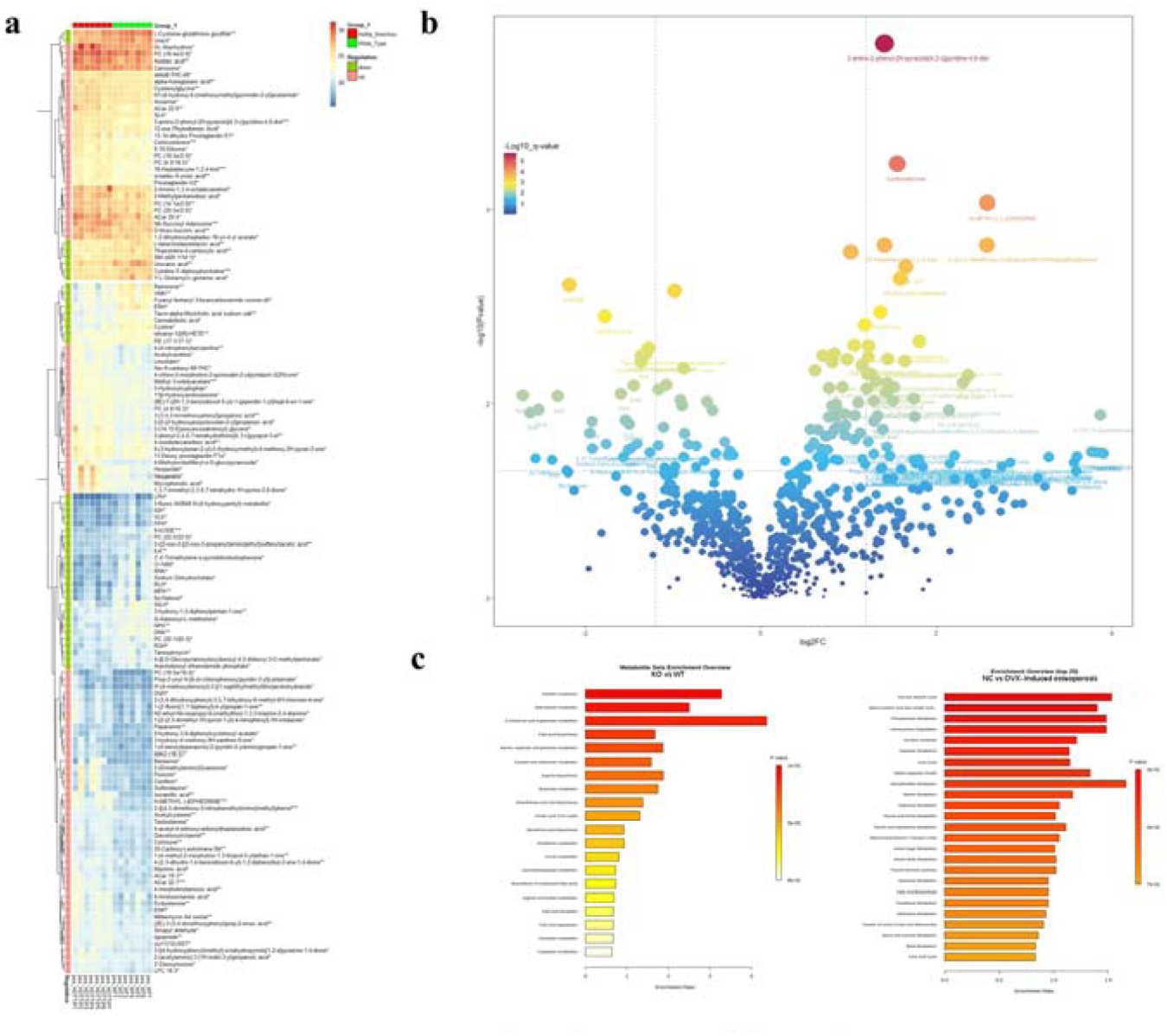

**Supplementary Figure 2.**
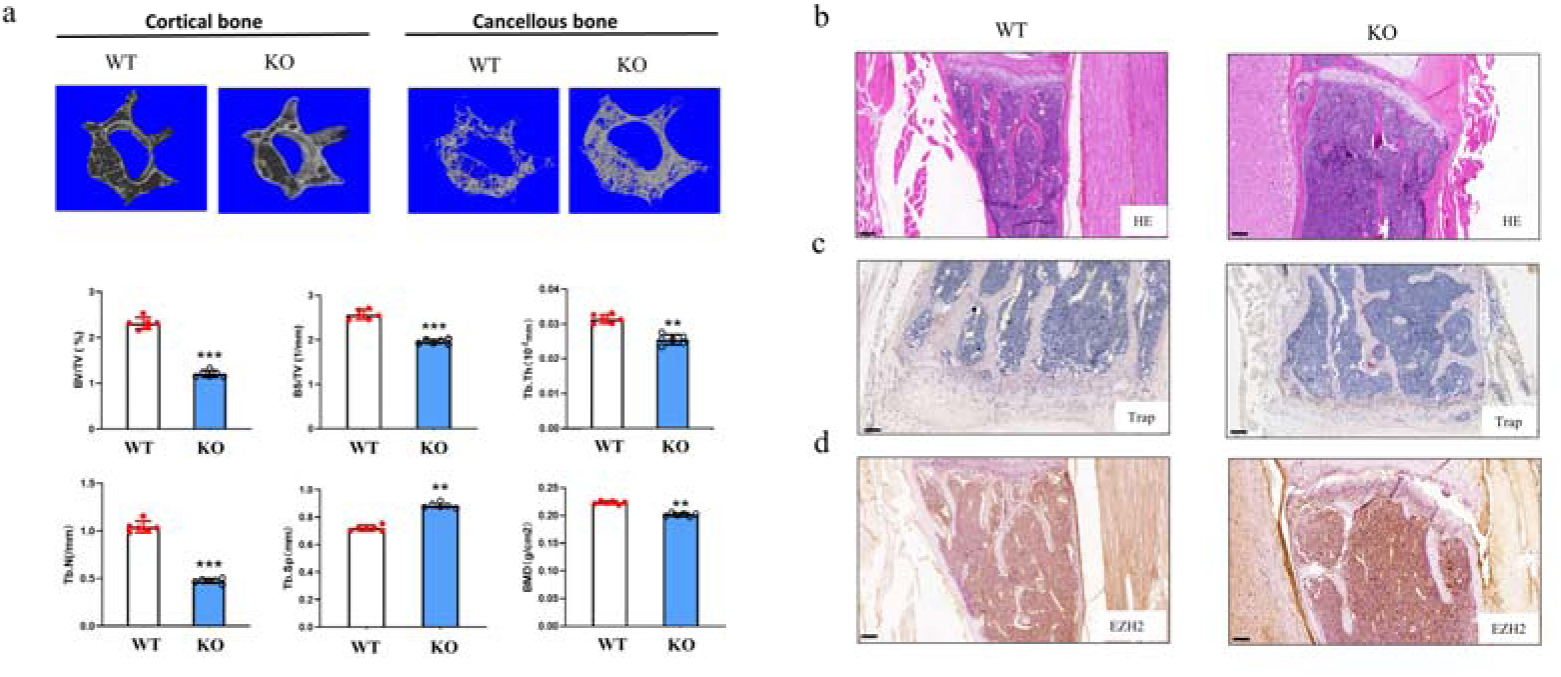

**Supplementary Figure 3.**
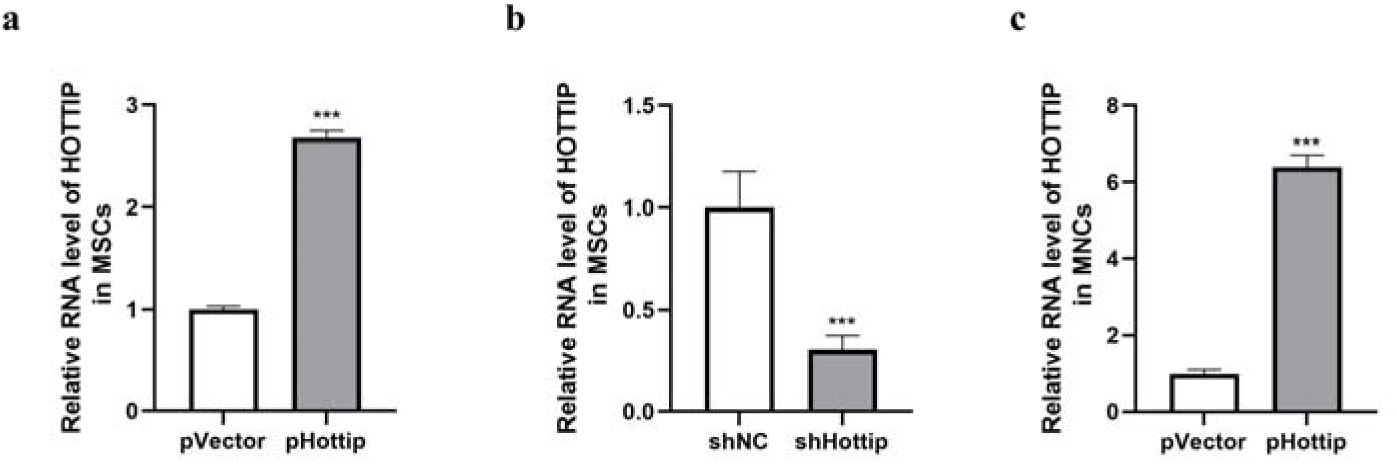

**Supplementary Figure 4.**
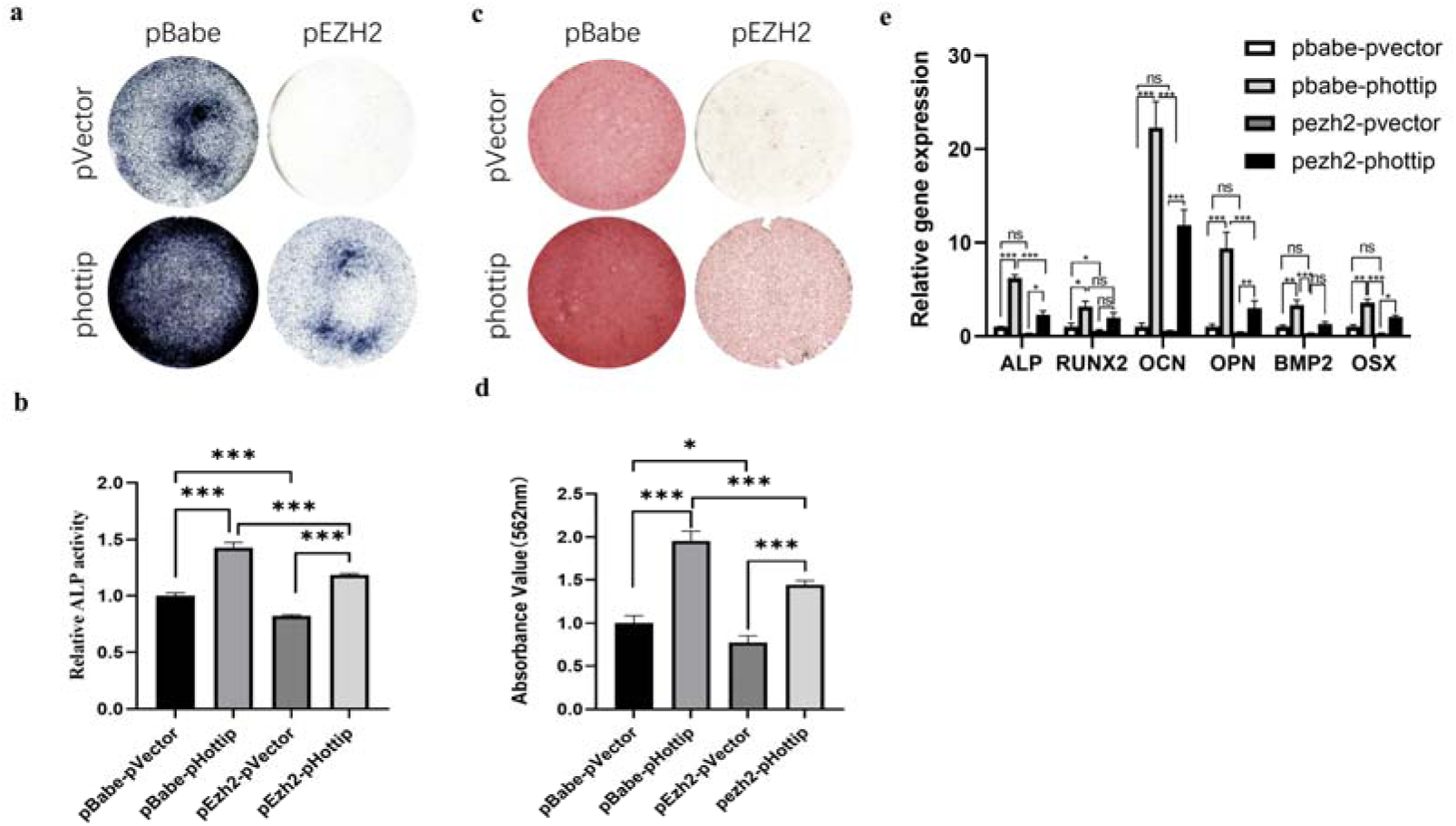

**Supplementary Figure 5.**
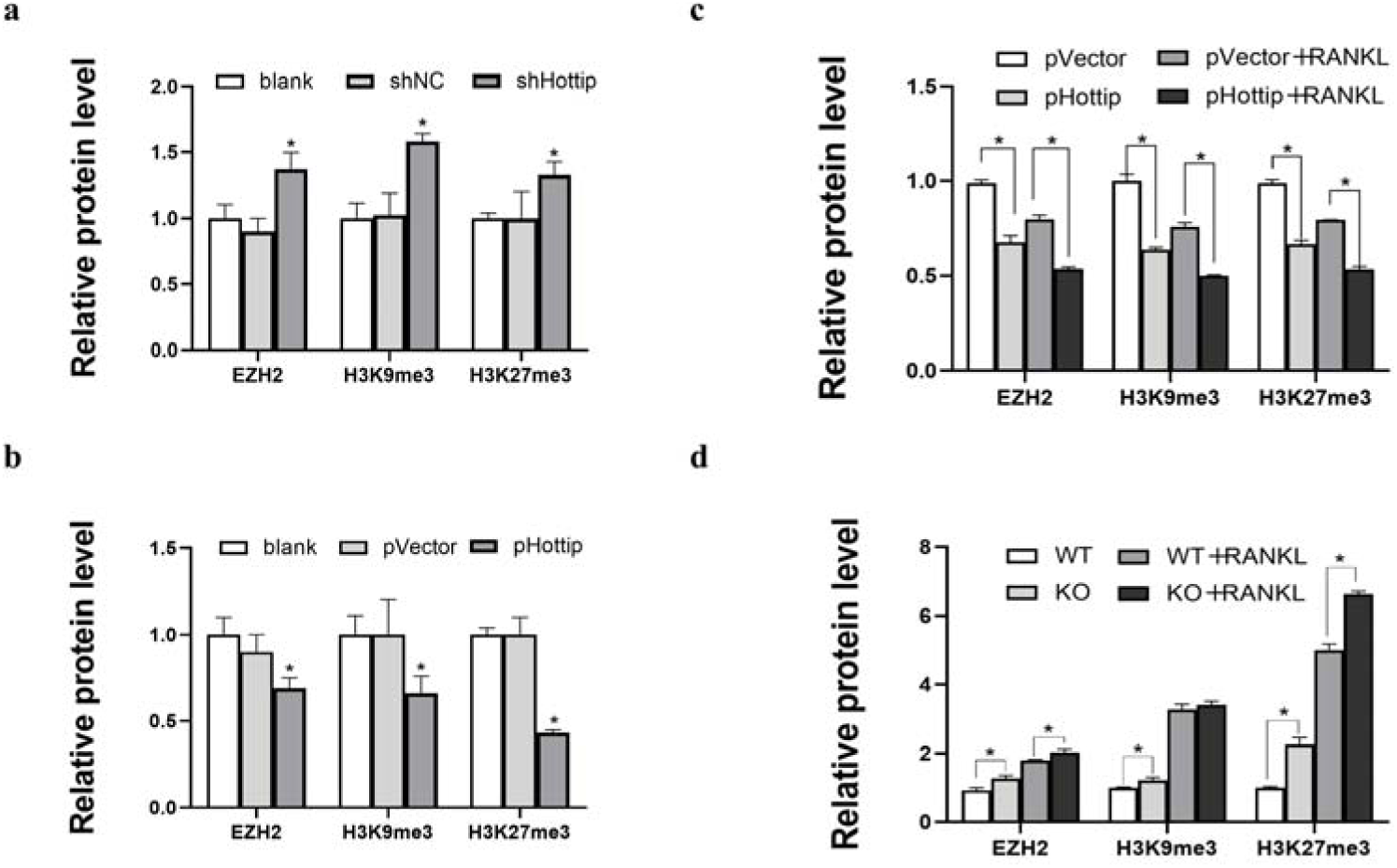

**Supplementary Figure 6.**
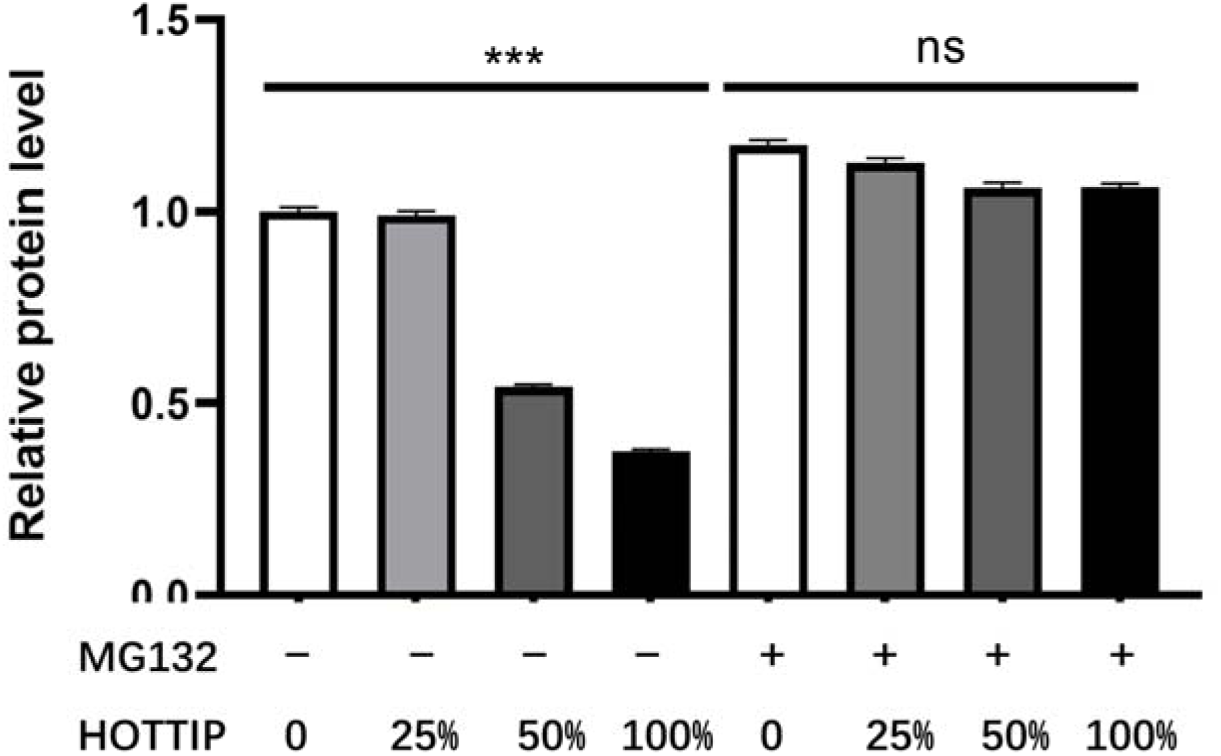

